## supplementary_materials for "The proportional recovery rule redux Arguments for its biological and predictive relevance"

### *Table of null values*

In the table below, we provide values of $\mathrm{cor}(x, \delta)$ for a range of input values $\mathrm{cor}(x, y)$, keeping $k$ fixed at 1. These are a result of equation (1) in the manuscript, and are intended to provide context for correlations between baseline and change that arise through coupling rather than as a result of a recovery process that affects the variance ratio.

| $k$ | $\mathrm{cor}(x, y)$ | $\mathrm{cor}(x, \delta)$ |
|---:|---:|---:|
| 1 | 0 | -0.707 |
| 1 | 0.1 | -0.671 |
| 1 | 0.2 | -0.632 |
| 1 | 0.3 | -0.592 |
| 1 | 0.4 | -0.548 |
| 1 | 0.5 | -0.5 |
| 1 | 0.6 | -0.447 |
| 1 | 0.7 | -0.387 |
| 1 | 0.8 | -0.316 |
| 1 | 0.9 | -0.224 |

### *Derivation of expressions for $\widehat{\beta_1}$ and $R^2$*

In the main paper, we discuss a simple linear regression of $\delta$ on ii $= \max - x$, where $\max$ is the maximum possible value of the scale and $\max - x$ is initial impairment. This regression can be written

$$\delta = \beta_0 + \beta_1 \text{ii} + \epsilon.$$

The inclusion of an intercept in the simple linear regression differs from the usual formulation of the PRR, but we find model helpful because it connects correlations to regression parameters and summaries. Specifically, the following expressions relate regression parameters and diagnostics to cor$(x, y)$ and $k$:

- $\widehat{\beta_1} = 1 - \text{cor}(x, y)\sqrt{k}$
- $R^2 = \text{cor}(x, \delta)^2 = \left(\frac{\widehat{\beta_1}}{\sqrt{k - 2\widehat{\beta_1} - 1}}\right)^2$

In a simple linear regression, the OLS estimate of the intercept is the ratio of the covariance of the predictor and response and the variance of the predictor. In this specific regression, we have:

$$\begin{aligned}
\widehat{\beta_1} &= \frac{cov(\delta, ii)}{var(ii)} \\
&= \frac{cov(y - x, \max - x)}{var(\max - x)} \\
&= \frac{cov(y, -x) + cov(-x, -x)}{var(x)} \\
&= \frac{var(x)}{var(x)} - \frac{cov(y, x)}{var(x)} \\
&= 1 - \frac{\text{cor}(x, y)\sigma_x \sigma_y}{var(x)} \\
&= 1 - \text{cor}(y, x)\frac{\sigma_y}{\sigma_x} \\
&= 1 - \text{cor}(y, x)\sqrt{k}
\end{aligned}$$

Next, we note that in a simple linear regression, $R^2$ is the squared correlation between the outcome and response. Then:

$$\begin{aligned}
R^2 &= \text{cor}(ii, \delta)^2 \\
&= \text{cor}(\max - x, \delta)^2 \\
&= \text{cor}(x, \delta)^2
\end{aligned}$$

Lastly, for linear regressions that include an intercept, $R^2$ is the squared correlation between the outcome and fitted values obtained from the model. Starting from this, we find:

$$
\begin{aligned}
R^2 &= \text{cor}\left(\widehat{\beta_0} + \widehat{\beta_1} ii, \delta\right)^2 \\
&= \left(\frac{cov\left(\widehat{\beta_0} + \widehat{\beta_1} ii, \delta\right)}{\sqrt{var\left(\widehat{\beta_0} + \widehat{\beta_1} ii\right) var(\delta)}}\right)^2 \\
&= \left(\frac{cov\left(\widehat{\beta_0} + \widehat{\beta_1}(max - x), y - x\right)}{\sqrt{var\left(\widehat{\beta_0} + \widehat{\beta_1}(max - x)\right) var(y - x)}}\right)^2 \\
&= \left(\frac{cov\left(-\widehat{\beta_1} x, y - x\right)}{\sqrt{var\left(-\widehat{\beta_1} x\right) var(y - x)}}\right)^2 \\
&= \left(\frac{cov\left(-\widehat{\beta_1} x, y - x\right)}{\sqrt{var\left(-\widehat{\beta_1} x\right) var(y - x)}}\right)^2 \\
&= \left(\frac{\widehat{\beta_1} cov(-x, y - x)}{\widehat{\beta_1}\sqrt{var(x) var(y - x)}}\right)^2 \\
&= \left(\frac{var(x) - cov(x, y)}{\sqrt{var(x)[var(x) + var(y) - 2cov(x, y)]}}\right)^2 \\
&= \left(\frac{var(x) - cor(x, y)\sqrt{var(x) var(y)}}{\sqrt{var(x)\left[var(x) + var(y) - 2cor(x, y)\sqrt{var(x) var(y)}\right]}}\right)^2 \\
&= \left(\frac{var(x) - cor(x, y) var(x)\sqrt{k}}{\sqrt{var(x)\left[var(x) + var(x)k - 2cor(x, y) var(x)\sqrt{k}\right]}}\right)^2 \\
&= \left(\frac{var(x)\left(1 - cor(x, y)\sqrt{k}\right)}{var(x)\sqrt{1 + k - 2cor(x, y)\sqrt{k}}}\right)^2 \\
&= \left(\frac{1 - cor(x, y)\sqrt{k}}{\sqrt{k + 2 - 2cor(x, y)\sqrt{k} - 1}}\right)^2 \\
&= \left(\frac{\widehat{\beta_1}}{\sqrt{k - 2\widehat{\beta_1} - 1}}\right)^2
\end{aligned}
$$

#### *Specification of the generalized additive model*

We use a generalized additive model (GAM) to allow smooth, non-linear associations between baseline $x$ and follow-up $y$. This is intended to provide a flexible candidate model for comparison with other mechanistic model implementations, including the PRR and constant recovery in the presence of strong ceiling effects.

Our implementation uses the `gam` function in the `mgcv` R package (Wood, 2012). A thorough overview of the theoretical and practical underpinnings of this widely-used package can be found in related monograph (Wood, 2017). Briefly, the default `gam` approach estimates non-linear associations using a rich thin-plate spline expansion with an explicit penalization to enforce smoothness in the result; for intuition, a high degree of penalization results in linear fits, while less penalization allows for greater non-linearity. The smoothing parameter(s) are selected using generalized cross validation as part of the model-fitting procedure.

Code supplements contain all model fitting procedures; for clarity, we note that our implementations are of the form:

```
mgcv::gam(y ~ s(x), data = winters_df)
```

### *Fitted values across training / testing splits*

We use cross validation to compared the performance of five models in terms of predictive. We considered an intercept-only model, a model assuming constant recovery with a ceiling effect, an additive model, and the PRR. For each dataset, we generated 1000 training / testing splits; results for prediction accuracy using MAPEs are shown in Figure 6 in the main paper. The Figure below shows fitted values for each model (except the intercept-only model) applied to training datasets generated in the cross validation procedure. The constant recovery model is visually a poor fit, and occasionally the additive model is too flexible, especially for the Zarahn data. The additive and exponential models yield fitted values that are similar to the PRR despite their additional flexibility and complexity.

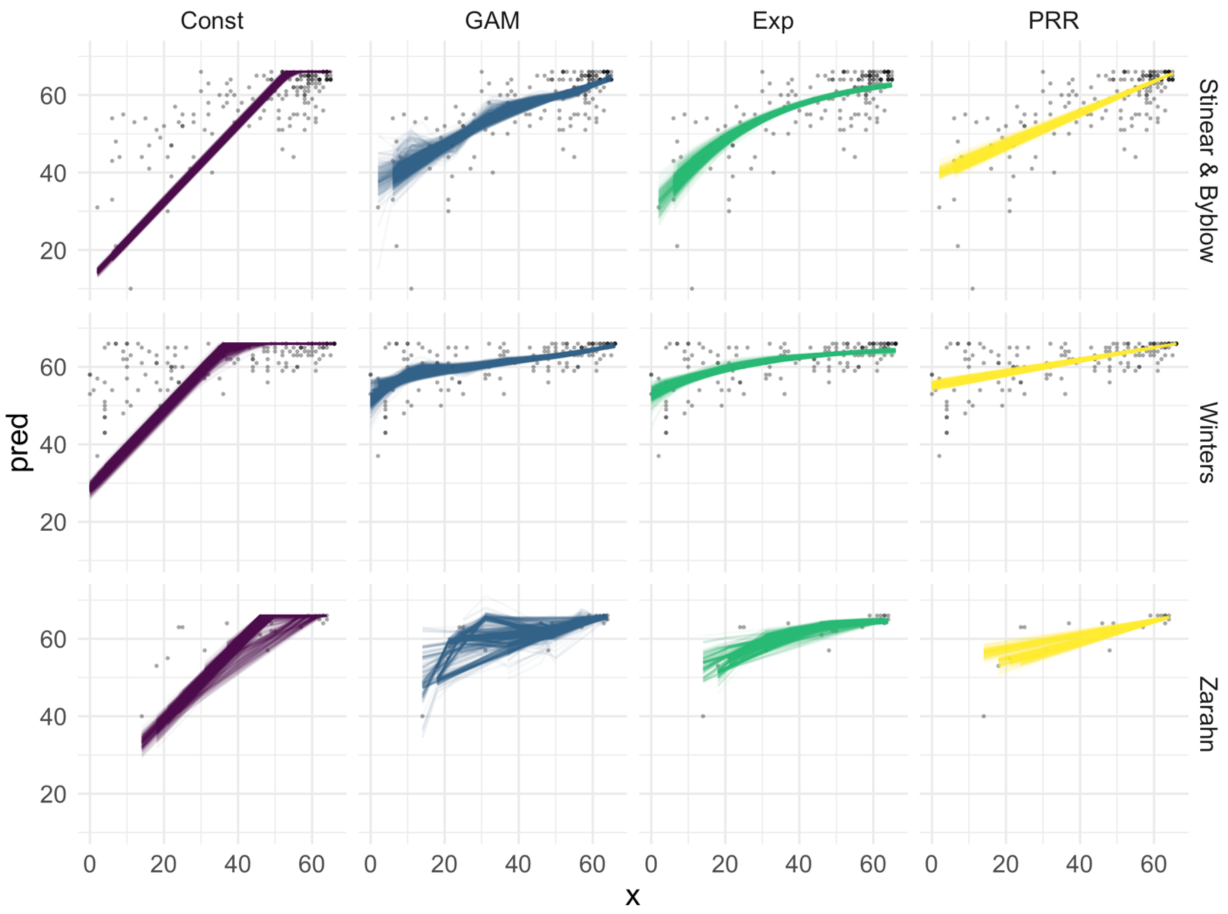
